# Store-operated Ca^2+^ entry regulatory factor (SARAF) alters murine metabolic state age-dependent via hypothalamic pathways

**DOI:** 10.1101/2022.08.03.500424

**Authors:** Diana Gataulin, Yael Kuperman, Michael Tsoory, Inbal E. Biton, Izhar Karbat, Anna Meshcheriakova, Eitan Reuveny

**Affiliations:** Department of Biomolecular Sciences, Weizmann Institute of Science, Rehovot 760001, Israel; Department of Veterinary Resources, Weizmann Institute of Science, Rehovot 760001, Israel

## Abstract

Store-operated Ca2+ entry (SOCE) is a vital process aimed at refilling cellular internal Ca^2+^ stores, and a primary cellular-signaling driver of transcription factors entry to the nucleus. SARAF (SOCE associated regulatory factor)/TMEM66 is an endoplasmic reticulum (ER) resident transmembrane protein that promotes SOCE inactivation and prevents Ca^2+^ overfilling of the cell. Here we demonstrate that mice deficient in SARAF develop age-dependent sarcopenic obesity with decreased energy expenditure, lean mass and locomotion without affecting food consumption. Moreover, SARAF ablation reduces hippocampal proliferation, modulates the activity of the hypothalamus-pituitary-adrenal (HPA) axis, and mediates changes in anxiety-related behaviors. Interestingly, selective SARAF ablation in the paraventricular nucleus (PVN) of the hypothalamus protects from old age-induced obesity and preserves locomotion, lean mass and energy expenditure, suggesting an opposing, site-specific role for SARAF. Lastly, SARAF ablation in hepatocytes leads to elevated SOCE, elevated vasopressin-induced Ca^2+^ oscillations, and an increased mitochondrial spare respiratory capacity, thus providing insights into the cellular mechanisms that may affect the global phenotypes. These effects may be mediated via the liver X receptor (LXR) and IL-1 signaling metabolic regulators explicitly altered in SARAF ablated cells. In short, our work supports both central and peripheral roles of SARAF in regulating metabolic, behavioral, and cellular responses.

**Highlights:**

- Loss/absence of SARAF facilitates age-dependent obesity with decreased metabolic rate, lean mass, and locomotion, without affecting food consumption.
- Loss of SARAF leads to lipid droplet hypertrophy, BAT whitening and age-dependent hepatic steatosis.
- Mice lacking SARAF expression in the PVN have an increased metabolic rate, decreased BAT whitening, and are protected from sarcopenic obesity.
- SARAF ablation in hepatocytes increases SOCE, elevates Ca^2+^ oscillation in response to vasopressin, and increases the mitochondria’s spare respiratory capacity.
- Loss of SARAF leads to decreased hippocampal proliferation, sensitized HPA-axis and changes in anxiety-related behavior.

## Introduction

Overweight and obesity affects almost 2 billion people worldwide (as calculated by World Health Organization-WHO in 2016) and is considered a 21st-century pandemic. Diet and exercise are considered the primary prevention and treatment strategies (https://www.who.int/health-topics/obesity#tab=tab_1). However, understanding the genetic factors involved in the central circuitry, metabolism, and adipose tissue homeostasis may improve the ability to treat obesity with precision (González-Muniesa et al., 2017; Van Der Klaauw and Farooqi, 2015). Metabolic homeostasis of the entire organism is a balanced feedback mechanism involving both the CNS and periphery, where the hypothalamic paraventricular nucleus (PVN) plays a central role (Kim et al., 2018);it directly and indirectly regulates critical hormones as corticosterone (cortisol in humans), adrenaline, vasopressin, oxytocin, thyrotropin-releasing hormone (TRH) (Herman et al., 2016). Of which regulate the body’s metabolic state (Qin et al., 2018a). Many intracellular pathways respond to these signaling cues to affect metabolic homeostasis, where changes in intracellular Ca^2+^ levels are central (Berridge, 2016; Clapham, 2007; Pozzan et al., 1994). It is thus critical to have tight regulation of intracellular and intraorganelle Ca^2+^ levels. This task is a well-coordinated action by many proteins that either pump Ca^2+^ from the cytosol or into intracellular organelles such as mitochondria and the endoplasmic reticulum (ER) (Pozzan et al., 1994).

Store-operated calcium entry (SOCE) is one of several processes that participate in the cell’s Ca^2+^ homeostasis (Prakriya and Lewis, 2015). In most cell types, it replenishes the Ca^2+^ cellular stores, like the ER and mitochondria, and in some it also plays a crucial role in transcription factor’s entry to the nucleus to activate gene transcription (Hogan et al., 2010). SOCE is activated following the release of Ca^2+^ from ER stores by inositol tri-phosphate (IP_3_), the product of the breakdown of phosphatidylinositol diphosphate (PIP2) by PLCβ or PLCγ activation via G_q_/11-coupled G-protein-linked receptors, or receptor tyrosine kinases, respectively (Berridge, 1993; Putney, 1986). Two essential components are required for SOCE activity, stromal interaction molecule (STIM), an ER transmembrane protein with a Ca^2+^ sensing domain at its ER luminal side, and a plasma membrane (PM) resident Ca^2+^ channel, (Orai) (Feske et al., 2006; Liou et al., 2005; Prakriya et al., 2006; Roos et al., 2005; Yeromin et al., 2006). SOCE is activated following the depletion of Ca^2+^from the ER lumen, oligomerization of STIM, and activation of Orai at PM-ER junctions to allow the flow of Ca^2+^ ions into the cell, to replenish the depleted stores. An additional layer of SOCE activity is provided by the interactions of STIM and Orai with the transient receptor potential channels (TRPCs), non-selective cation channels (Ambudkar et al., 2016; Liao et al., 2007).

Calcium signaling involving the ER and mitochondria is compromised in obesity and metabolic diseases (Arruda and Hotamisligil, 2015; Berridge, 2016). SOCE is a crucial element in lipid metabolism and adiposity; specifically, it plays a role in the mobilization of fatty acids from lipid droplets, lipolysis, and mitochondrial fatty acid oxidation (Baumbach et al., 2014; Cuk et al., 2017). Additionally, SOCE is necessary for glucose-stimulated pancreatic insulin secretion (Sabourin et al., 2015; Tamarina et al., 2005). Moreover, Ca^2+^ signaling and SOCE are involved in proper liver function, including bile secretion, proliferation, oscillatory response to hormones, cholesterol, and glucose metabolism (Amaya and Nathanson, 2013; Aromataris et al., 2008). Alterations in liver SOCE result in decreased ER Ca^2+^ content, ER stress, inflammation, impaired insulin function, and abnormal glucose metabolism (Arruda et al., 2014; Wilson et al., 2015). Furthermore, SOCE was found to be impaired in the liver of obese murine models (Arruda et al., 2017; Park et al., 2010). SOCE was also implicated in myocytes’ function, specifically its importance in the proper function of the sarcoplasmic reticulum (Stiber et al., 2008; Wei-Lapierre et al., 2013; Zhao et al., 2015). Furthermore, SOCE function in myocytes distinguishes between aged and young muscle (Thornton et al., 2011; Zhao et al., 2008).

SARAF is an ER resident protein that we previously reported to associate with STIM and promote SOCE inactivation (Palty et al., 2012). SARAF plays a key role in shaping cytosolic Ca^2+^ signals and determining the content of the major intracellular Ca^2+^ stores, a role that is probably important in protecting the cell from Ca^2+^ overfilling. SARAF is localized to PIP2-rich PM junctions (Cao et al., 2015) and is activated by dimerization at its luminal end (Kimberlin et al., 2019) when stores are full, by a still unknown mechanism. Reduced SARAF levels increase intracellular SOCE and was recently found to be involved in several pathological conditions such as pancreatitis and cerebral ischemia (La Russa et al., 2020; Son et al., 2019). Moreover, SARAF is an androgen-responsive marker for prostate cancer and it regulates mTOR-dependent cardiac growth (Romanuik et al., 2009; Sanlialp et al., 2020). Here we report on the consequences of knocking out SARAF, both globally and in PVN-specific manner in mice. KO of SARAF introduces a new role for SARAF in physiological context. SARAF KO mice gain weight and loose lean mass at a later age. This weight gain is not the consequence of increased food intake but dependent on PVN-associated circuits. At the cellular level, SARAF increases mitochondrial respiration in addition to its expected reduction of SOCE activity. This report may place SARAF as an important component in the pathophysiology of obesity.

## Results and Discussion

### Generation of SARAF-KO mice

To study its physiological role, we used the knock-out mouse project (KOMP) repository, first to create SARAF LacZ knock-in cassette inserted mice and then SARAF loxP-floxed conditional mice, termed SARAF^fl/fl^, as described in the experimental procedures section and graphical representation (Figure 1A). SARAF was previously shown to be highly expressed in the immune and neuronal tissues (Palty et al., 2012); we focused on its expression in the brain via LacZ staining and found it has a marked expression in the hippocampus, hypothalamus (specifically the PVN) and the amygdala (Figure 1B). Later, by crossing the conditional SARAF^fl/fl^ mice to the ubiquitously expressing PGK promoter-driven Cre recombinase mice to generate PGK-Cre^+^:SARAF^fl/fl^, whole - body knock-out mice, termed SARAF-KO. The knockout of SARAF was validated via Western blotting of the brain tissue and compared with their littermates SARAF^fl/fl^PGK-Cre^-^, who were homozygous to the floxed SARAF, but did not express Cre recombinase, termed SARAF-WT (Supp. 1A). SARAF knock-out mice were viable and bred normally with Mendelian distribution.

**Figure 1:**
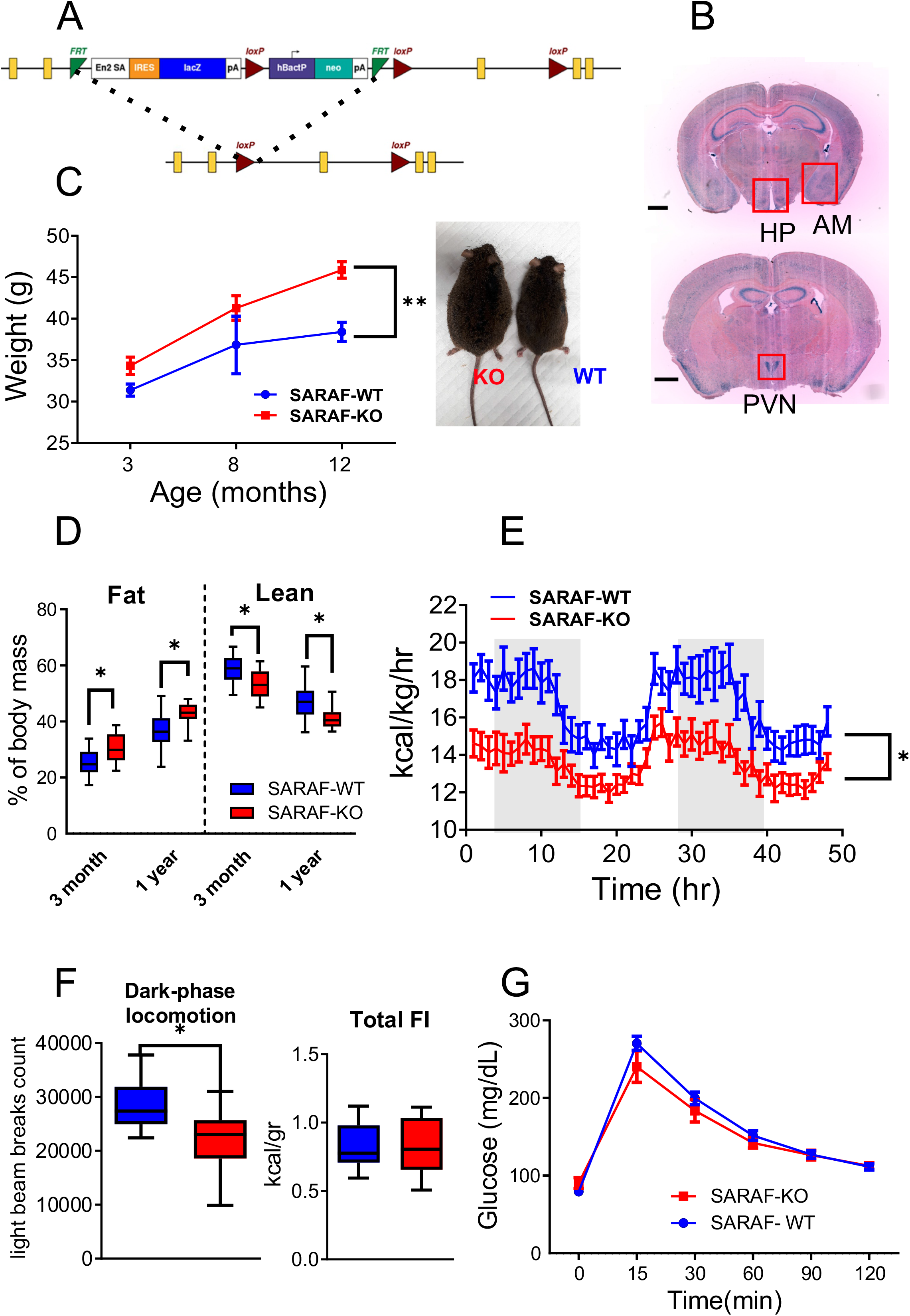
Generation of SARAF conditional mice, SARAF expression in the brain and SARAF^fl/fl^ PGK-Cre^+^ metabolic phenotype. **A,** Schematic representation of the SARAF knock-in cassette and after flipase exertion of the cassette leaving crucial exon 3 flanked by loxP sites. **B**, X-gal-stained coronal brain sections of KOMP cassette-inserted heterozygous mice, expressing β-gal at the sites of SARAF expression (scale bar 1mm). **C**, Changes in body weight of SARAF-WT (n=13, blue) and SARAF-KO (n=11, red) over time. Inset, representative photos of SARAF-WT (right) and SARAF-KO (left) mice at 1-years-old. **D**, Lean and fat mass percentage of SARAF-WT (n=13, blue) and SARAF-KO (n=11, red) mice at 3-months and 1 years old. **E-F**, PhenoMaster calorimetry metabolic analysis of 3-month-old mice (SARAF-WT, n=8; SARAF-KO, n=10). E. Heat production over time. **F,** Dark-phase locomotion. **G**, Total food intake. **H,** three-months old mice glucose tolerance test ((SARAF-WT, n=7; SARAF-KO, n=9).

### SARAF-KO mice have impaired metabolic function and lipid accumulation

Male SARAF-KO mice were compared with their WT littermates and were found to have a significant increase in body weight. The weight differences were significantly higher from early adulthood (three months), where the KO mice weighted on average about 10% more than the WT mice. The weight differences increased with age. At 1 year old KO mice weighted nearly 20% more than the WT mice (Figure 1C). Linear growth was not altered in the SARAF-KO mice (Supp.1B). Interestingly, the weight differences were also reflected in body composition alterations, including an increased percentage of fat mass and lower percentage of lean mass in the KO mice, suggesting a possible role for SARAF in mediating sarcopenic obesity-like symptoms (Figure 1D). Indirect calorimetry assessment of SARAF-KO mice revealed a lower metabolic rate (as manifested in heat production) and reduced locomotion (measured during the active phase in the diurnal cycle) (Figure 1E-F). The abovementioned changes were not associated with significant differences in food intake (Supp.1C). Moreover, food intake after five hours fasting in a refeed experiment did not alter either the weight loss or the weight gain following the experiment (Supp.1D). Interestingly, despite the altered body weight and the body composition, the mice were not diabetic or pre-diabetic since they present with a normal fasting glucose level and had a normal response to a glucose load (as assessed by glucose tolerance test (GTT)) (Figure 1G).

We then examined the adipose tissue distribution in the elderly SARAF-KO mice using computed tomography (CT) imaging and found that the excess fat was distributed throughout the body with a high tendency for abdominal accumulation (Figure 2A-C). Hematoxylin and eosin (H&E) stained white adipose tissue (WAT) droplets size analysis of inguinal and visceral adipose tissues (iWAT and vWAT, respectively), revealed significant hypertrophy of fat cells. Moreover, we found that fat was accumulated both in the intracapsular brown adipose tissue (iBAT) and in the liver, resulting in iBAT whitening and hepatic steatosis (Figure 2D-H). These histological phenotypes were seen in 3-month-old mice and upsurge in 1-year-old mice, except liver fat accumulation, which is not obvious yet in younger mice (Supp.1E-I). BAT is an organ that contributes to systemic metabolic homeostasis and thermoregulation, and its size is associated with various pathologic conditions including obesity (Enerbäck, 2010; Shimizu and Walsh, 2015). Interestingly, iBAT whitening can be prevented by inhibiting Ca^2+^ overload in the mitochondria, (Gao et al., 2020). Moreover, the hepatic steatosis is accompanied by cellular Ca^2+^ imbalance and is a predisposition for a non-alcoholic fatty liver disease that might lead to liver cirrhosis and cancer (Ali et al., 2017). The weigh- and fat-associated changes in elderly SARAF ablated mice were also accompanied by subclinical hypothyroidism, manifested by normal serum thyroxine and cholesterol, and elevated TSH levels (Fatourechi, 2009)(Figure 2I-K). Subclinical elevation of TSH might influence resting metabolic rate and may hint at early thyroid dysfunction (Tagliaferri et al., 2001). The correlation between the old age increase in TSH and hepatic steatosis may raise the possibility of a thyroid-liver interaction (Huang et al., 2013).

**Figure 2:**
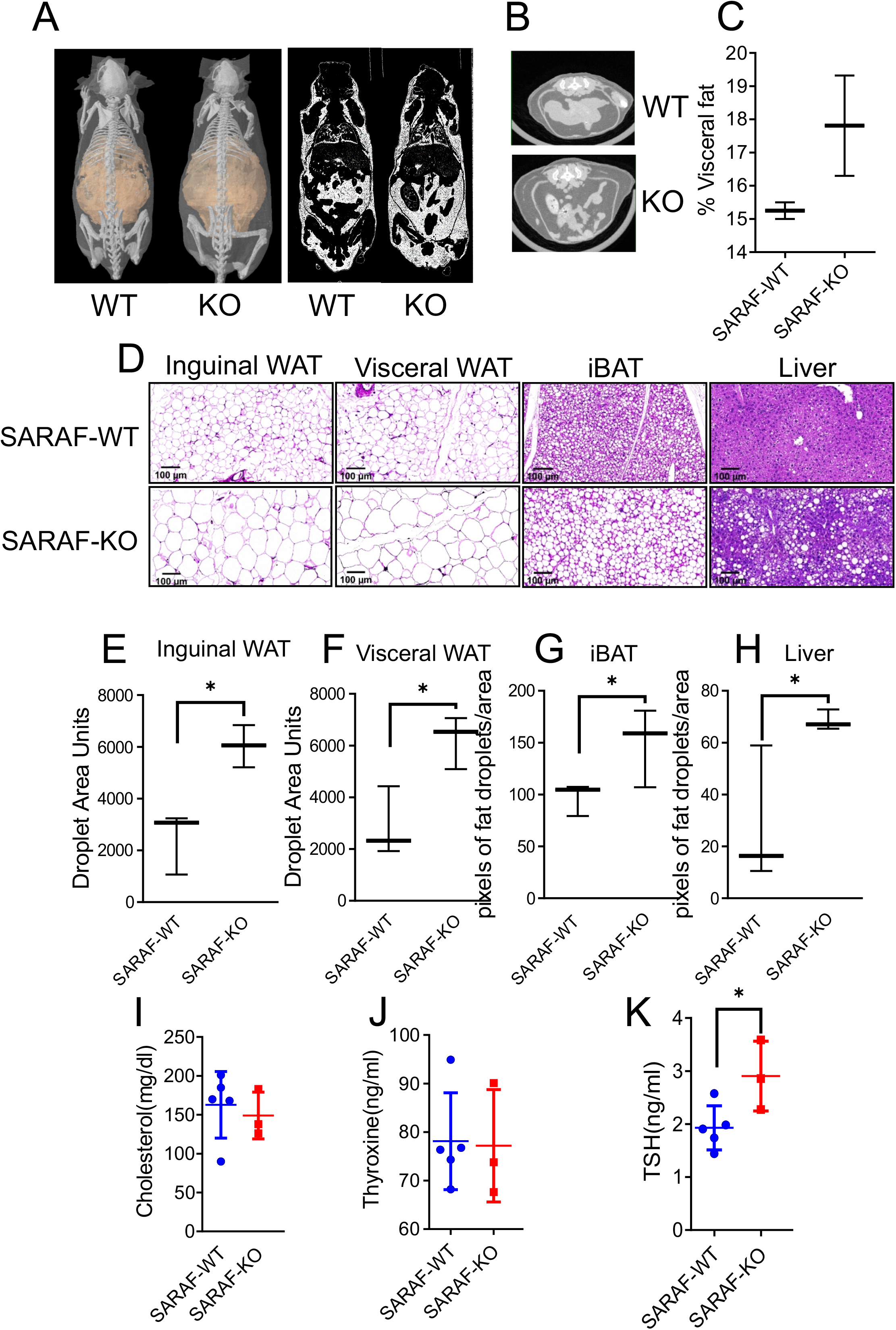
Characterization of SARAF-KO lipid deposition phenotype and voluntary training treatment. **A,** Micro-CT imaging and whole-body fat (right) distribution analysis of representative 1-year-old SARAF-WT and SARAF-KO mice. **B**, Abdominal fat representative images of CT images of above-mentioned mice. **C**, Quantification of visceral fat deposition. **D**, Inguinal WAT, visceral WAT, liver and iBAT tissue histology in 1-years-old SARAF-WT and SARAF-KO mice. Size bar-100 μm. **E-F**, Inguinal, and visceral WAT droplet size quantification. **G-H**, iBAT, and liver pixels of fat droplets/area quantification. **I,** Serum analysis of cholesterol levels of 1 year-old mice. **J**, Serum analysis of thyroxine levels 1 year-old mice. **K**, Serum analysis of TSH levels of 1 year-old mice.

### SARAF ablation in SIM1 neurons improves metabolic function and hints toward hypothalamic metabolic feedback

SARAF regulated metabolic phenotypes discussed above could stem from any several metabolic organs, including adipose, muscle and brain derived regulation. Because of the marked SARAF expression in the hypothalamus, we chose to focus on the role of hypothalamic SARAF in energy homeostasis. To this end, we crossed the SARAF^fl/fl^ mice to SIM1 promoter-driven Cre-recombinase expressing mice (SIM1-Cre) (Balthasar et al., 2005) to induce SARAF deletion specifically in the hypothalamic PVN neurons, termed SARAF-SIM1KO. Mice were validated via site-specific punch-PCR of the PVN (Suppl. 2A). Unlike SARAF-KO, three month old SARAF-SIM1KO mice had similar body weights and composition to their WT littermates. However, at the age of one year old, these mice were found to have improved age-related metabolic phenotypes, inversely mirroring the whole body SARAF-KO. At 1 year old, SARAF-SIM1KO mice had lower body weight with a reduced percentage of fat mass and an increased percentage of lean mass than SARAF-SIM1WT littermates (Figure 3A-B). Like the whole body knocked-out mice, SARAF-SIM1KO mice did not have an altered response to glucose, or differences in food consumption either under basal or refeed challenged conditions (Figure 3C and Supp. 2B-C). Moreover, their heat production and locomotor activity were elevated in the opposite tendency to the SARAF-KO mice (Figure 3D-F). Those phenotypes are clear only at old age (over 1-years-old) except for increased locomotion, which was already evident in 3-month-old mice (Supp.2D-2F). The increased locomotion in SARAF-SIM1KO mice is of interest since it is consistent from a young age to old age and thus may drive the phenotype by increasing lean mass and/or by changing energy expenditure. Several studies have implicated hypothalamic PVN in the regulation of locomotion (Gladfelter and Brobeck, 1962; McCormick and Njeri Ibrahim, 2006; Mönnikes et al., 1992). Interestingly, when examining lipid accumulation in the SARAF-SIM1KO mice, we noticed that BAT had reduced fat accumulation compared with the wild type littermates. We did not witness differences in WAT droplet size or fat accumulation in the liver (Figure 3G-K). When challenging these mice with diet rich in fat and carbohydrates food (Western diet) for 18 weeks, their total increases in body weight, composition and calorimetry analysis, were similar between the two groups (Supp.3A-E). In Western diet fed mice lipid accumulation phenotype, glucose tolerance, and refeed responses were unchanged as well (Supp. 3E-J), thus strengthening the assumption that SARAF related metabolic phenotypes in SARAF SIM1KO mice are not related to food intake related mechanism. These results point toward the coordination of BAT and thermogenesis by the PVN via the Ca^2+^ homeostasis mechanism (Qin et al., 2018b). It is of note that SARAF SIM1KO mice did not exhibit altered anxiety-related behavioral phenotype suggesting that SARAF, although ubiquitously expressed, plays a site-specific role (supp. 4G-I).

**Figure 3:**
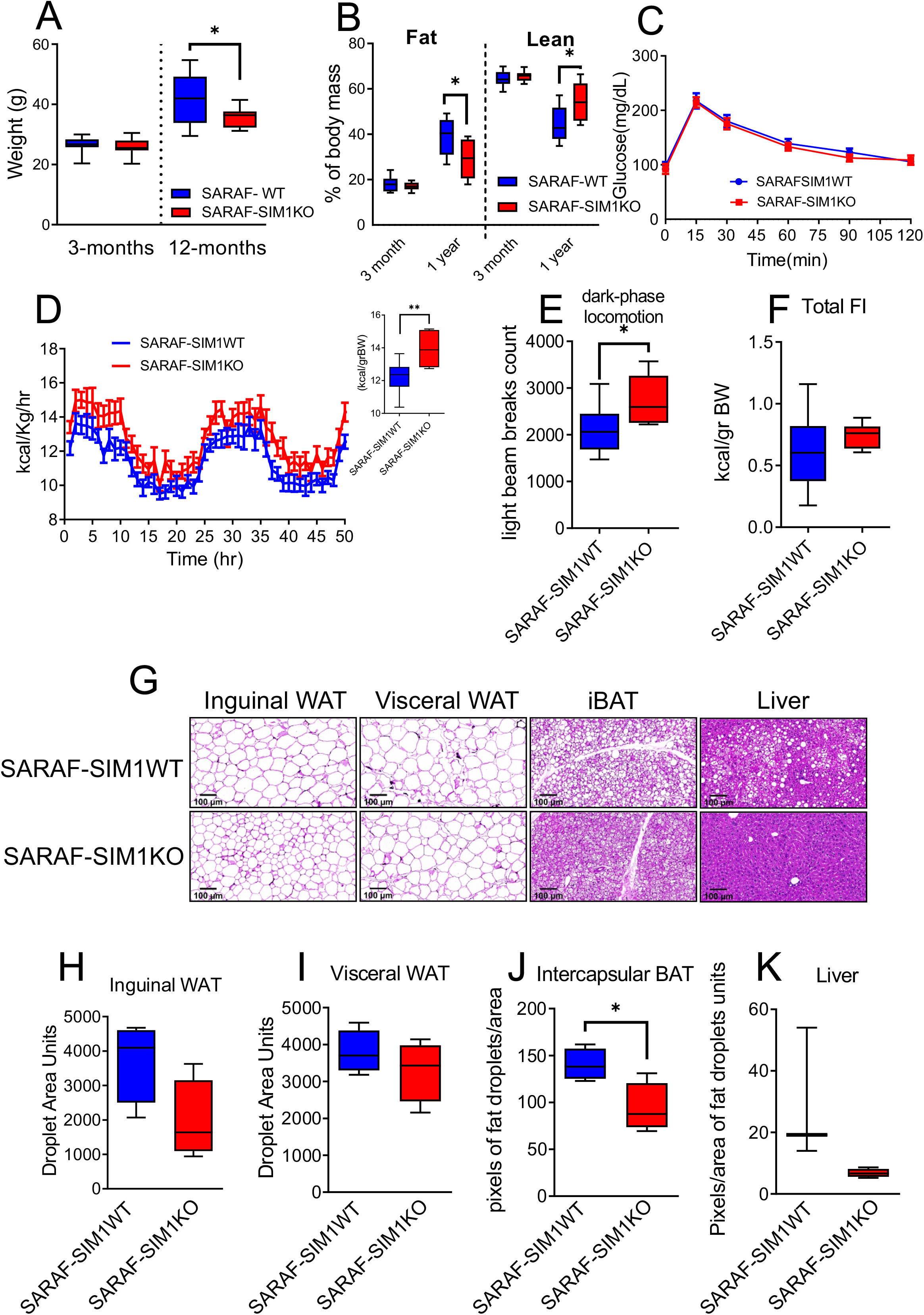
SARAF-SIM1KO mice metabolic related phenotypes. **A**, three-months, and 1-year old SARAF-SIM1WT (n=16) and SARAF-SIM1KO (n=9) mice body weight comparison. **B**, Lean and fat mass percentage SARAF-SIM1WT and SARAF-SIM1KO three-months and one-year old mice of the same mice as in A. **C**, Three-months-old glucose tolerance test (SARAF-SIM1WT, n=11; SARAF-SIM1KO, n=6). **D-F**, PhenoMaster calorimetry metabolic analysis of one-year old mice (SARAF-SIM1WT, n=10; SARAF-SIM1KO, n=6). **D**, Heat production over time, and heat production per body weight (inset). **E**, Dark-phase locomotion. **F**, Total food intake. **G**, Inguinal WAT, visceral WAT, liver and iBAT tissue histology in one-years-old SARAF-SIM1WT and SARAF-SIM1KO mice. Size bar-100μm. **H-I**, Inguinal, and visceral WAT droplet size quantification. **J-K**, iBAT and liver pixels of fat droplets/area quantification.

### SARAF ablation decreases hippocampal proliferation, affects the stress response and HPA-axis activation

SARAF was highly expressed in the hippocampus (Figure 1B) and specifically in a subset of doublecortin (DCX) positive neuronal progenitors, indicative of proliferating neuronal stem cells (Supp.4A). Since hippocampal proliferation has a marked impact on anxiety and metabolism via the HPA-axis (Kanoski and Grill, 2017; Snyder et al., 2011), we sought to examine the impact of SARAF on hippocampal proliferation and its effect on the hypothalamus-pituitary-adrenal (HPA) axis, as, altered function of this system may affect the metabolic phenotype and may account, in part, for the phenotypes reported above. EdU (5-ethynyl-2’-deoxyuridine) incorporation was used to examine hippocampal cell proliferation in the SARAF-WT and in the SARAF-KO mice. EDU incorporation assay revealed a significant decrease in proliferation in the dorsal and the ventral hippocampus dentate gyrus in the SARAF-KO mice (Figure 4A). This decrease was independently validated via an immunostaining against proliferating cell nuclear antigen (PCNA) as the proliferation marker (supp. 4B) and repeated in primary mouse embryonic fibroblast (MEF) cultures derived from SARAF-WT and SARAF-KO using staining for Ki67 expression (Supp. 4C).

**Figure 4:**
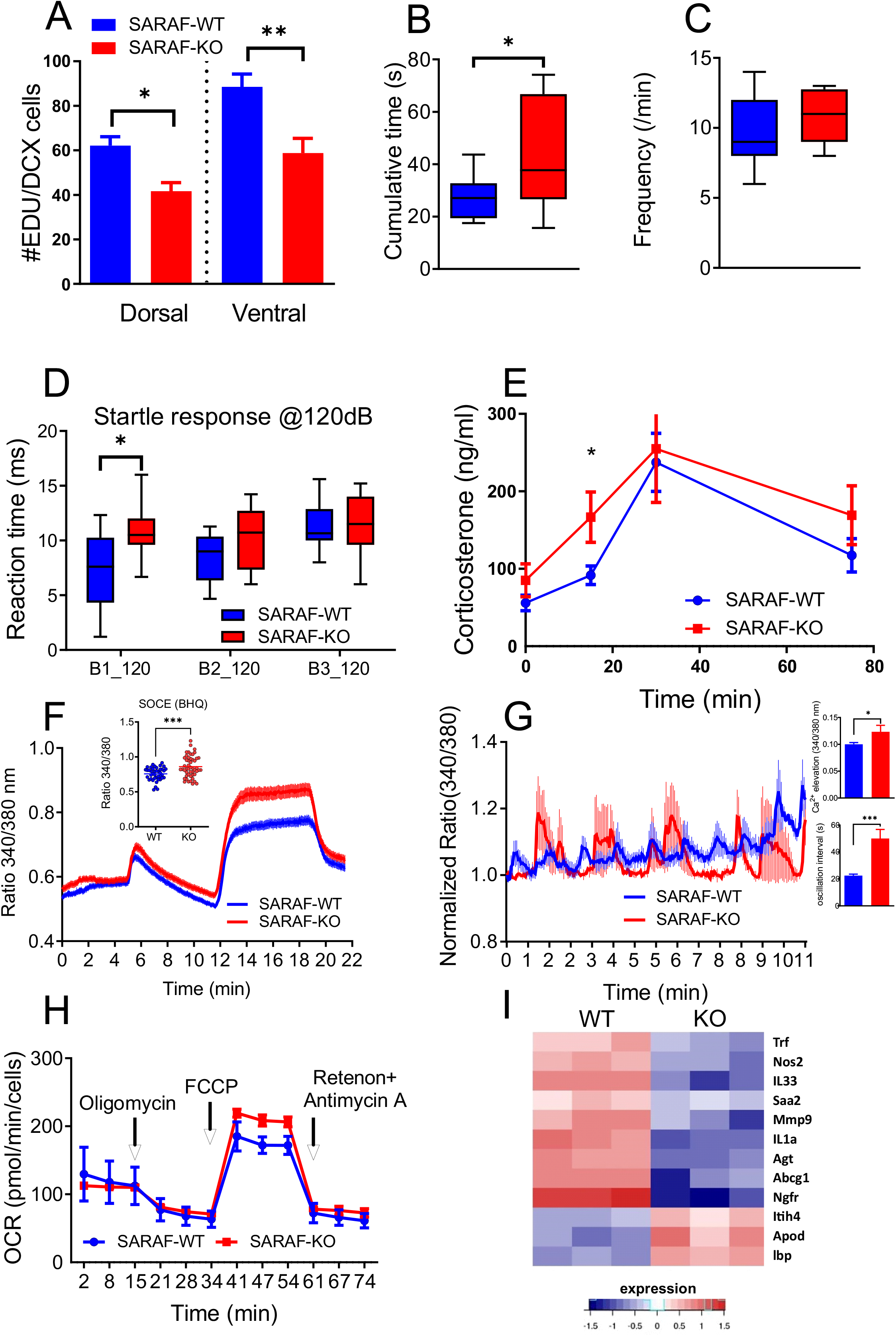
Hippocampal neurons proliferation, anxiety related behavioral phenotypes and hepatocyte cellular phenotypes of SARAF^fl/fl^PGK-Cre^+^ mice. **A**, EDU(5-ethynyl-2’-deoxyuridine) proliferation analysis of SARAF-WT and SARAF-KO mice at the dorsal and ventral hippocampal dentate gyrus proliferation quantification. **B-C**, 3-month-old SARAF-WT (n=11) and SARAF-KO (n=10) mice performance in anxiety-related tests: **B**, dark-light transfer. **C**, Frequency of exits. **D**, Acoustic startle response test. Reaction time in 3 blocks of 120db stimuli. **E**, Blood corticosterone levels following 15-minute restrain stress (SARAF-WT, n=8; SARAF-KO, n=7). **F**, Fura-2AM Ca^2+^ imaging trace of primary hepatocytes extracted from SARAF-WT and SARAF-KO mice and the quantification of SOCE levels (inset). **G**, Ca^2+^ oscillations in hepatocytes induced by vasopressin (1nM) as measured by FURA-2AM. Quantification of Ca^2+^elevation and oscillation interval (inset). **H**, Mitochondrial respiration of SARAF-WT and SARAF-KO primary hepatocytes and their spare respiratory capacity. **I**, Heatmap representation of LXR/RXR hepatic (liver X receptor) pathway activation which was indicated by ingenuity platform in MEF RNA sequencing results.

Behaviorally, SARAF-KO mice exhibited decreased anxiety-like behavior as manifested in the dark-light transfer (DLT) by increased time spent in the lit compartment (Figure 4B-C), and in the acoustic startle response (ASR) experiment, where they exhibited longer reaction time in the first set of stimulus presentations (Figure 4D). Counterintuitively, SARAF knocked out mice exhibited a mild increase in HPA-axis activation, as suggested by a stronger immediate corticosterone response to restraint-stress (Figure 4E). This discrepancy may stem from differences in the timing of assessment; 5 to 10 min following the initiation of the exposure to stress in the DLT and ASR tests, as opposed to about 30 min in the CORT assessment. Interestingly, despite the reduced neurogenesis observed, long-term memory, assessed using the Morris water maze (Supp. 4F), and short-term memory, assessed using the Y-maze (Supp. 4D-E), were not altered.

### SARAF ablation leads to increased cellular SOCE, elevated mitochondrial spare respiratory capacity, and altered gene expression patterns

Cellular metabolic functions influence global metabolic phenotypes and behavior; specifically, mitochondrial respiration greatly influences global metabolic phenotypes (Picard et al., 2015). Moreover, since the hypothalamus, directly via neuronal connections and indirectly via hormonal secretion, regulate metabolic organs (Gore, 2013; Roh et al., 2016), we sought to examine whether the cellular metabolic functions were consistent with the global phenotypes we observed in the KO animals. Specifically, the liver is tightly regulated by the PVN via a direct neuronal innervation and hormonal regulation (O’Hare and Zsombok, 2016; Uyama et al., 2004). For this reason, we extracted primary hepatocytes from SARAF-KO mice and examined their Ca^2+^ signaling and SOCE by FURA-2AM-based Ca^2+^ imaging. We induced SOCE by the transient sarco/endoplasmic reticulum Ca^2+^-ATPase (SERCA) inhibitor BHQ. SOCE activity was elevated in the knocked-out hepatocytes, confirming our previous in vitro experiments (Palty et al., 2012). Hepatocyte SOCE was inhibited by the Orai inhibitors La^3+^ and 2-ABP, therefore, hinting at the involvement of the classical SOCE and the involvement of the STIM-Orai machinery (Bird and Putney, 2018) (Figure 4F).

Ca2+ oscillations are a significant driver of cellular signaling, and it are mediated by SOCE and the mitochondrial Ca^2+^ uptake (Dolmetsch et al., 1998). Vasopressin is a pituitary secreted hormone that regulates several physiological functions, including behavior, thermoregulation, water absorption, liver function, and adipogenesis (Jones et al., 2008; Maus et al., 2017). When examining Ca^2+^ oscillations triggered by physiological levels of vasopressin (1nM), we observed a marked increase in release amplitude and increased oscillation intervals in the knocked-out hepatocytes (Figure 4G). The latter strengthens the idea that SARAF may affect hypothalamic control over metabolic organs, including the liver. Next, we sought to examine cellular metabolic function by measuring mitochondrial respiration by directly examining the cellular oxygen consumption rate (Traba et al., 2016). This function is highly influenced by the cellular Ca^2+^ levels and influences global metabolic phenotypes (Böhm et al., 2020; Gherardi et al., 2020; Picard et al., 2015). Cellular mitochondrial respiration was markedly altered in SARAF knocked out hepatocytes, having a significantly higher spare respiratory capacity (SRC) function (Figure 4H). These findings suggest that SARAF, via the modulation of SOCE activity, has a great influence on metabolic organs at a cellular level, like the liver, with altered mitochondrial metabolism and SOCE. Since Ca^2+^, in addition to its direct regulatory action, can affect at longer time scales gene transcription, we set to examine gene transcription patterns by RNA sequencing embryonic fibroblast (MEF) cells derived from SARAF-WT and SARAF-KO animals. We analyzed the results for canonical pathway enrichment using QIAGEN’s Ingenuity^®^ software and identified that LXR (Liver X receptor)/RXR (Retinoid X receptor) signaling pathway was significantly altered (Figure 4I), hinting, again, at SARAF importance in proper liver function. The LXR/RXR signaling pathway genes that were altered include IL1α, IL33, LBP, and ApoD, all of which have indications in metabolic function (Jakobsson et al., 2012; Miller et al., 2010). LXR/RXR and the related LXR regulated IL-1-signaling pathways were previously shown to influence metabolism via the hypothalamus and directly influencing adipose tissue and the liver (Barbier et al., 2019; Ericsson et al., 1994; Hong and Tontonoz, 2014; Kruse et al., 2017).

## Conclusion

This study demonstrates that SARAF is involved in the physiological regulation of age-dependent metabolic rate (as manifested by energy expenditure) and locomotion, without affecting food consumption or clear blood glucose handling. Moreover, SARAF was involved in a complex metabolic phenotype that includes lipid metabolism regulation in the adipose tissue, the BAT and liver. These global phenotypes were accompanied by SARAF-mediated cellular metabolic functions, including hepatocyte’s SOCE, vasopressin evoked Ca^2+^ oscillations and mitochondria’s spare respiratory capacity. Examining SARAF’s contribution to the central nervous system regulation functions, indicted its involvement in PVN regulated autonomic-sympathetic control of energy expenditure, locomotion, and BAT profile. Moreover, SARAF was found to regulate hippocampal proliferation, while having some effects on the HPA-axis and anxiety-related behavior.

Cellular Ca^2+^ balance is a major factor in fine-tuning metabolic function in different organs, including the liver, muscle, adipose tissues and neurons; even slight changes in Ca^2+^ levels might affect their function (Ong et al., 2019; Toescu and Verkhratsky, 2007). SARAF is expressed in the Sim-1 expressing neurons of the brain, the muscle and BAT. SOCE homeostasis, regulated by SARAF, is especially important in a subset of specific cells/tissues, described in this study, it is suggested, however, that additional subtle and yet to be discovered SARAF related physiological functions are probably affected as well. Moreover, other proteins essential for maintaining Ca^2+^ homeostasis, like SERCA, whose activity is reduced with aging (Lompré et al., 1991) may be more essential in the absence of SARAF. In this study, the influence of SARAF on SRC was indicated in the hepatocytes OCR measurements, underlining the importance of maintaining appropriate Ca^2+^ levels for healthy-cell function (Chemaly et al., 2018). Interestingly, SRC is a known influencer over aging muscle function, and a driver of sarcopenia (Hiona et al., 2010). Thus, SRC might be the driver of SARAF involvement in lean mass maintenance. However, in the liver, SARAF causes improved SRC, seemingly opposite to the previously described papers, however, increase in SRC is possible, in case of damaged oxidative phosphorylation as indicated in some reports (Porter et al., 2015). Interestingly, and relating to our running wheel data, training increases in aged muscle (Waters et al., 2003). Furthermore, SRC is also elevated in the hyperglycemic adipocytes, allowing for fast adaptation and recovery, SARAF ablation in the metabolic organ forces increasing SRC levels as an adaptive response to less than favorable conditions (Keuper et al., 2014).

SARAF role in hippocampal neurons is still in need of further refining; specifically, it is perplexing that SARAF involvement in hippocampal proliferation did not alter learning and memory. Nevertheless, it has been previously discussed that significant cellular aberration should occur to influence a behavioral phenotype (Dauer and Przedborski, 2003), given the importance of Ca^2+^ regulation for cellular physiology, it is possible that compensatory mechanisms exist to provide an additional layer of protection from an aberrant Ca^2+^ steady-state levels. This can be in the factors associated with SOCE, pumps that clear cytosolic Ca^2+^, or an increase in the cell-buffering capacity (Carreras-Sureda et al., 2013; Feng et al., 2006, 2010; Karakus et al.; Srivats et al., 2016).

Collectively, our study shows for the first time, to the best of our knowledge, that SARAF is an essential contributor to the dysregulation of general and PVN regulated metabolic states. Moreover, it is a novel model for age-dependent sarcopenic obesity, which is not dependent on feeding or is accompanied by diabetes. Pharmacological targeting of the SARAF signaling pathway may provide a novel approach for treating sarcopenic obesity.

## Acknowledgments

This study was supported, in part, by the US-Israel Binational Foundation (2015298), the Minerva Foundation, the Willner Family Fund and Yeda-Sela Center, all to E.R. E.R. is the incumbent of the Charles H. Hollenberg Professorial Chair.

## Materials and methods

### Mice

The SARAF conditional KO strain used for this research project was generated from KOMP ES cell line *saraf^tm1a(KOMP)Wtsi^*, RRID:MMRRC_061775-UCD, was obtained from the Mutant Mouse Resource and Research Center (MMRRC) at University of California at Davis, an NIH-funded strain repository, and was donated to the MMRRC by The KOMP Repository (University of California, Davis); Originating from Pieter de Jong, Kent Lloyd, William Skarnes, Allan Bradley, Wellcome Sanger Institute. The ES cells were created by introducing a splice acceptor/reporter cassette containing a poly-A site into an endogenous intron upstream of a critical exon. The cassette includes the lacZ gene and the neomycin resistance gene surrounded by FLP sites, and the critical (third) SARAF exon floxed by loxP sites (Fig. 1A). The following Jackson laboratory mice lines were used for crossing and generation of conditional mice lines: Gt(ROSA)26Sortm1(FLP1)Dym, Tg(Pgk1-cre)1Lni, B6.FVB(129X1) Tg(Sim1-cre)1Lowl/J. All general and spatially specific SARAF knocked-out mice were compared to their wild-type littermates.

All the animal procedures were approved by the Weizmann Institute Institutional Animal Care and Use Committee (IACUC).

### Genotyping

Genomic DNA was isolated from a tail biopsy using 25mM NaOH and 0.2mM disodium EDTA extraction buffer (PH12) incubated for 1 h at 95°C, the extract was neutralized using 40mM Tris-HCl neutralization buffer (PH5), and PCR identified mouse genotypes. Two PCR primers were synthesized to detect the intact *Cre* gene; three PCR primers were synthesized to detect the KOMP, WT, and loxp inserted SARAF gene, and four PCR primers were used to detect WT, KO, and HET mice in the SARAF^fl/fl^PGKCre line. The PCR conditions were 95°C denaturing, 60°C (for SARAF) 55°C (for Cre) 65°C (for SARAF^fl/fl^PGKCre) annealing, and 72°C extension for 35 cycles using Taq polymerase and a DNA Thermal Cycler from Bio-Rad. The PCR products were resolved by electrophoresis in 1% agarose gel.

### Mouse embryonic fibroblasts (MEFs)

E14.5 embryos were dissected, and their trunks were extracted into cold PBS. The trunks were finely minced using a razor until they were pipettable. Cells were suspended in 3ml trypsin-EDTA 0.05%, at 37C for 10 min. Cells were flushed through 21g (3ml) syringe three times, and double serum levels to block the trypsin activity. Next, cells were centrifuged at 1800 rpm for 5 min, and the pellet re-suspended in 8 ml of MEF media, containing 10% fetal calf serum, 2% Penstrep, 1% L-glutamine, and 1% sodium pyruvate. Cells from each embryo were plated on a 100mm dish coated with 0.1% gelatin. Cells were split 1:4 and frozen when they reached confluence (Parameswaran et al., 2008).

### Primary hepatocyte

Mouse hepatocytes were isolated from 8–12 weeks old male mice. Mice were anesthetized using ketamine and xylazine, liver was exposed and perfused by two steps, wash and a digestion step was done by Liberase research-grade (Roche Diagnostics). Cells were centrifuged in Percoll (GE Healthcare) gradient and seeded on Collagen type 1 (Sigma) coated plates with HBM basal medium+ HCM single Quotes (Lonza) media containing 1% FBS for 2.5–3 h, later washed with non-serum containing media and used for experiments for up to 24 hours (Halpern et al., 2018).

### Primary hippocampal cultures

Cultures were prepared from E18-P0 mouse pups, pups were decapitated, their brains removed, and the hippocampi dissected free and placed in chilled (4°C), oxygenated Leibovitz L15 medium (Gibco, Gaithersburg, MD, USA). The hippocampi were dissociated mechanically, and cells were plated in 24-well plate, onto round coverslips coated with L-polylysine. Glia plated 14 days beforehand and grown in 10%FBS and Glutamate. Cultures grew in a medium containing 5% HS, 5% FBS and B27 in an incubator at 37 °C with 5% CO2. Cells were imaged at 13–17 days after plating.

### Behavioral assays

All behavioral assays were performed during the dark period (08:00–20:00) in a reverse cycle room. Before each experiment, mice were habituated to the test room for two hours.

#### Morris water maze

For the acquisition phase, mice were subjected to 4 trials per day with an interval of 15 mins, for 7 consecutive days. In each trial, the mice were required to find a hidden platform located 1 cm below the water surface in a 120 cm-diameter circular pool. In the testing room, only distal visual-spatial cues for locating the hidden platform were available. The escape latency in each trial was recorded up to 90s. Each mouse was allowed to remain on the platform for 15s and was then removed from the maze. If the mouse did not find the platform within the 90s, it was manually placed on it for 15s. Memory was assessed 24 hours after the last trial. The escape platform was removed, and mice were allowed to search for it for 1 minute, and the time spent in the different quadrants of the pool was recorded using a VideoMot2 automated tracking system (Tse-System)(Panayotis et al., 2018; Ruggiero et al., 2017).

#### Y-maze

The maze contains three arms at 120 degrees to each other. The mouse underwent training and test on the same day. During training, one arm was closed. During the test, the mouse starts at the end of one arm and then chooses between the other two arms. The amount of time and duration spent in the closed arm is measured to demonstrate learning and memory (Maron et al.).

#### Dark-light transfer test

The test consists of a polyvinyl chloride box divided into a dark black compartment (14 × 27 × 26 cm) and a white 1050-lx illuminated light compartment (30 × 27 × 26 cm) connected by a small passage. Mice were placed in the dark compartment to initiate a 5-min test session. The time spent in the light compartment and the number of entries to the light compartment were measured.

#### Acoustic startle response

Mice were placed in a small Plexiglas mesh cage on top of a vibration-sensitive platform in a sound-attenuated, ventilated chamber. A high-precision sensor integrated into the measuring platform detected the movement. Two high-frequency loudspeakers inside the chamber produced all the audio stimuli. The acoustic startle response session began with 5 min acclimatization to white background noise [65 dB] maintained throughout the session. Thirty-two startle stimuli [120 dB, 40ms duration with a randomly varying ITI of 12–30s] were presented. The reaction time and latency to peak startle amplitude were measured (Panayotis et al., 2018).

### Immunohistochemistry

The mice were anesthetized and perfused with 1% PFA in PBS. Brains were carefully removed and fixed overnight in 30% sucrose 1% PFA in PBS. The following day 30 μm coronal slices were prepared using a sliding microtome and stored in PBS 0.01% Na-Azid.

For X-gal staining, the 30 μm slices were incubated in PBS containing 1% glutaraldehyde for 4 min and washed three times in PBS. Slices were incubated overnight in X-gal staining solution containing 3 mM K_3_Fe(CN)_6_, 3 mM K_4_Fe(CN)_6_, 1.3 mM MgCl_2_, 0.02% NPO_4_, 0.01% NaDOC and 1 mg/ml X-gal in PBS, filtered through a 0.45 μM filter. The next day slices were washed three times in PBS and mounted on slides.

For immunohistochemistry, the following primary antibodies were used: rabbit anti-beta-galactosidase (1:1000; Cappel), goat anti-doublecortin (1:100; Santa Cruz). Slices/cells were first incubated for 1.5 h at room temperature in a blocking solution containing 20% normal horse serum (NHS) and 0.3% triton x100 in PBS, followed by 48–72 h incubation at 4°C in primary antibody solution containing 2% NHS and 0.3% triton x100 in PBS. The slices were then washed with PBS and incubated with biotin-conjugated goat anti-rabbit (1:200; Jackson) in 2% NHS-PBS for 1.5 h at room temperature. Finally, slices were incubated for 1 h at room temperature with secondary antibody FITS -conjugated donkey anti-goat (1:200; Jackson) and Cy3-streptavidin (1:200; Jackson). Slices were then washed with PBS and incubated for 10min with Hoechst (1:2000), washed and mounted for fluorescence imaging.

Cells were fixed with 4% PFA in PBS for 15 min, washed, and permeabilized using 0.2% triton x100 in PBS for 30 min. Cells were blocked for 1.5 h, with 5% horse serum in PBS. The primary rabbit anti Ki67 (1:200, Abcam) antibody was incubated overnight. Secondary goat anti-rabbit cy3 (1:200, Jackson) antibody was incubated for 30 min. Cells were then washed with PBS and incubated for 10min with Hoechst (1:2000), washed and mounted for fluorescence imaging.

<u>EdU-proliferating cells staining</u> mice were given 0.2 mg/ml EdU (CarboSynth) in drinking water for two weeks and then sacrificed. Brains were embedded in paraffin, and hippocampal slices were then used for immunostaining, using the Click-iT EdU imaging kit (Invitrogen). Staining was done according to manufacturer instructions.

### Rabbit polyclonal anti-SARAF antibody production

Rabbit polyclonal anti SARAF antibody production was performed with the help of the Weizmann institute core facilities, antibody engineering unit. Two NZW SPF rabbits (New Zealand White rabbits) were subcutaneously injected with SARAF’s luminal domain (aa 30–164) fused to 6xHistidine tag (Kimberlin et al., 2019). The first injection was done with Freund’s complete adjuvant (Difco 263810), and the second was given a week later with Freund’s incomplete adjuvant (263910). Three boosters with an interval of 14 days were given IP in PBS, and serum was collected after each boost. The serums were purified for mouse IgG subclasses by affinity chromatography on protein-A Sepharose beads CL-4B (SPA-Sepharose, Pharmacia, Sweden). The best serum of the two was selected (according to western blotting), and additional purification steps were conducted as follows: rabbit serum was filtered using a 0.2mM filter (Nalgene) and loaded onto the affinity HisTrap-NHS column. The column was washed by 10CV PBS, 15CV washing buffer, and an additional 10CV PBS. The first elution step was performed with a Glycine buffer (0.1M Glycine titrated to pH 2.3 with HCl). The second elution step was then done using the 5CV of TEA (Tetra-Ethyl-Ammonium) pH 11.5 (titrated with NaOH). The second fraction, eluted under basic conditions, had superior selectivity and specificity compared to the total serum and wad used for western blot analysis. All columns used are commercially available as pre-packed media from GE healthcare.

### Western blot

Protein was extracted from adult mice brains in sucrose extraction buffer containing 0.32 M sucrose, 2mM EDTA, 10mM HEPES pH=7.4, PMSF (1:100), and protease inhibitor cocktail (1:50). Extracts were then homogenized on ice with glass homogenizer, centrifuged at 750g for 10 min to remove crude debris, then ultracentrifuged at 200,000 g for 40 min, pellet re-suspended in CL47a complexiolyte solubilization buffer (Logopharm) and kept for 30 min on ice and then used for protein quantification with Pierce BCA kit (Thermo Fisher). Protein lysates were electrophoresed on a 10% SDS–polyacrylamide gel electrophoresis (SDS-PAGE), transferred to nitrocellulose membranes, blocked and treated overnight with a rabbit polyclonal anti-SARAF (1:5000), in PBS containing 0.05% tween (PBST) with 1% BSA. After washing, the membranes were incubated with horseradish peroxidase-conjugated goat anti-rabbit IgG antibody (Jackson) in PBST with 2.5% skim milk. Electrochemiluminescence (ECL) detected the immunoreactive protein (WesternBright ECL HRP substrate, Advansta).

### Corticosteroid blood measurements

Corticosterone was measured 5 h after the dark cycle begun using the DetectX Corticosterone CLIA kit (Arbor assays). 5μl tail blood samples from mice were collected before (basal), immediately after 15 min of restraint stress, 30 and 75 min from stress initiation. The restraint stress was induced using a cut 50-ml plastic conical tube. Plasma samples were immediately centrifuged and stored at −80 °C until assays for hormone measurement were conducted. Blood was analyzed according to manufacture instructions (Lebow et al.).

### micro-CT

Mice were anesthetized with isoflurane (3% for induction, 1%–2% for maintenance) mixed with oxygen (1 l/min) and delivered through a nasal mask. Once anesthetized, the mice were placed in a head-holder to assure reproducible positioning inside the scanner. The set of mice were scanned using a micro-CT device TomoScope® 30S Duo scanner (CT Imaging, Germany) equipped with two source-detector systems. The operation voltages of both tubes were 40 kV. The integration time of protocols was 90 ms (360 rotation) for 3 cm length, and axial images were obtained at an isotropic resolution of 80 μ. Due to the maximum length limit, to cover the entire mouse body, imaging was performed in two parts with the overlapping area, and then all slices merged to one dataset representing the entire ROI. The radiation dose for each mouse was 2.2 Gy(Akselrod-Ballin et al., 2016). Fat quantification analysis was performed using a CT analysis (skyscan Bruker MicroCT) software (Version1.19).

### Calcium Imaging

Cells were plated onto 24 mm L-polylysine coated cover glass 4-24 h before the experiment. Before the experiment, the cover glass was mounted on an imaging chamber and washed with a 0/2mM Ca^2+^ solution. Fura-2AM loading of cells was performed for 30-45 minutes. Cytosolic Ca^2+^ levels were recorded from Fura-2AM-loaded cells, excited at wavelengths of 340/20 and 380/20 nm and imaged with 510/80 nm filters. For all single-cell imaging experiments, traces of averaged responses, recorded from 10 to 50 cells.

Cells were stimulated with ATP (100mM), carbachol (100mM) and bradykinin (1.5mM) in 100mg/ml BSA or with S-DHPG (10mM)/Vasopressin(1nM) and SOCE was induced by 2-5mM of Ca^2+^. Additionally, cells were washed with 0mM Ca^2+^ and stores were emptied by thapsigargin (2mM)/BHQ (20mM) and SOCE was again induced by 2-5mM Ca^2+^. All materials were purchased from Sigma and diluted in 0/2mM Ca^2+^ solution. Solutions used for imaging were either standard Ringer’s solution with or without CaCl2 or solutions with the following constituents: 0mM Ca^2+^ solution contained HBSS-/-, 20mM Hepes, 1mM MgCl_2_, 0.5mM EGTA and glucose calibrated to PH=7. 2-5 mM Ca^2+^ solution contained the same except for the absence of EGTA and addition of 2-5 mM CaCl_2_(Kimberlin et al., 2019; Palty et al., 2012).

### Murine metabolic studies

Indirect calorimetry, food, water intake, and locomotor activity were measured using the LabMaster system (TSE-Systems, Bad Homburg, Germany). Data were collected after 48h of adaptation from singly housed mice. Body composition was assessed using the Bruker minispec mq7.5 live mice analyzer (Wolf et al., 2017).

### Running wheel

Mice were singly housed in standard cages equipped with a running wheel for four weeks (Colombus Instruments). Distances were recorded every 15 min from a counter attached to the wheel. The wheel circumference (111.76 cm) was converted to kilometers.

### Cellular respiration

Measurement of intact cellular respiration was performed using the Seahorse XF24 analyzer (Seahorse Bioscience Inc.) and the XF Cell Mito Stress Test Kit according to the manufacturer’s instructions. Respiration was measured under basal conditions, and in response to Oligomycin (ATP synthase inhibitor; 0.5 μM) and the electron transport chain accelerator ionophore, FCCP (trifluorocarbonylcyanide phenylhydrazone; 1 μM), to measure the maximal oxygen consumption rate (OCR). Finally, respiration was stopped by adding the electron transport chain inhibitor AntimycinA (1 μM) (Ruggiero et al., 2017).

### Glucose tolerance test

Mice were fasted for five hours and subsequently given 2g/kg glucose solution by i.p. injection. Blood glucose was determined at 0, 15, 30, 60, 90, and 120 min after the glucose challenge (FreeStyle Freedom Lite, Abbott)(Wolf et al., 2017).

### RNA sequencing

Total RNA was extracted from the indicated cell cultures using the RNeasy kit (QIAGEN). Then, RNA integrity was evaluated on a Bioanalyzer (Agilent 2100 Bioanalyzer), requiring a minimal RNA integrity number (RIN) of 8.5. Libraries were prepared according to Illumina’s instructions accompanying the TruSeq RNA Sample Preparation Kit v2 (cat # RS-122–2001).

According to the manufacturer’s instructions, sequencing was carried out on Illumina HiSeq 2500v4 SR60, 20 million reads per sample.

Sequenced reads were mapped to the Mus musculus genome version GRCm38, using TopHat v2.0.10. Genes were identified using a.gtf obtained from Ensembl release 82. Per gene, reads were counted using HTSeq. Normalization of read counts and P-values for differentially expressed genes were computed using DESeq2(Kalisky et al., 2018).

### Statistical and image analysis

Images were analyzed and quantified using the Fiji/ImageJ software.

P-values were calculated using a student’s t-test for statistical comparisons of mean values using the GraphPad Prism software. Differences were regarded significant for p<0.05 (*) and highly significant for p<0.01 (**) and p<0.001(***).

## STAR methods

### Key resources table

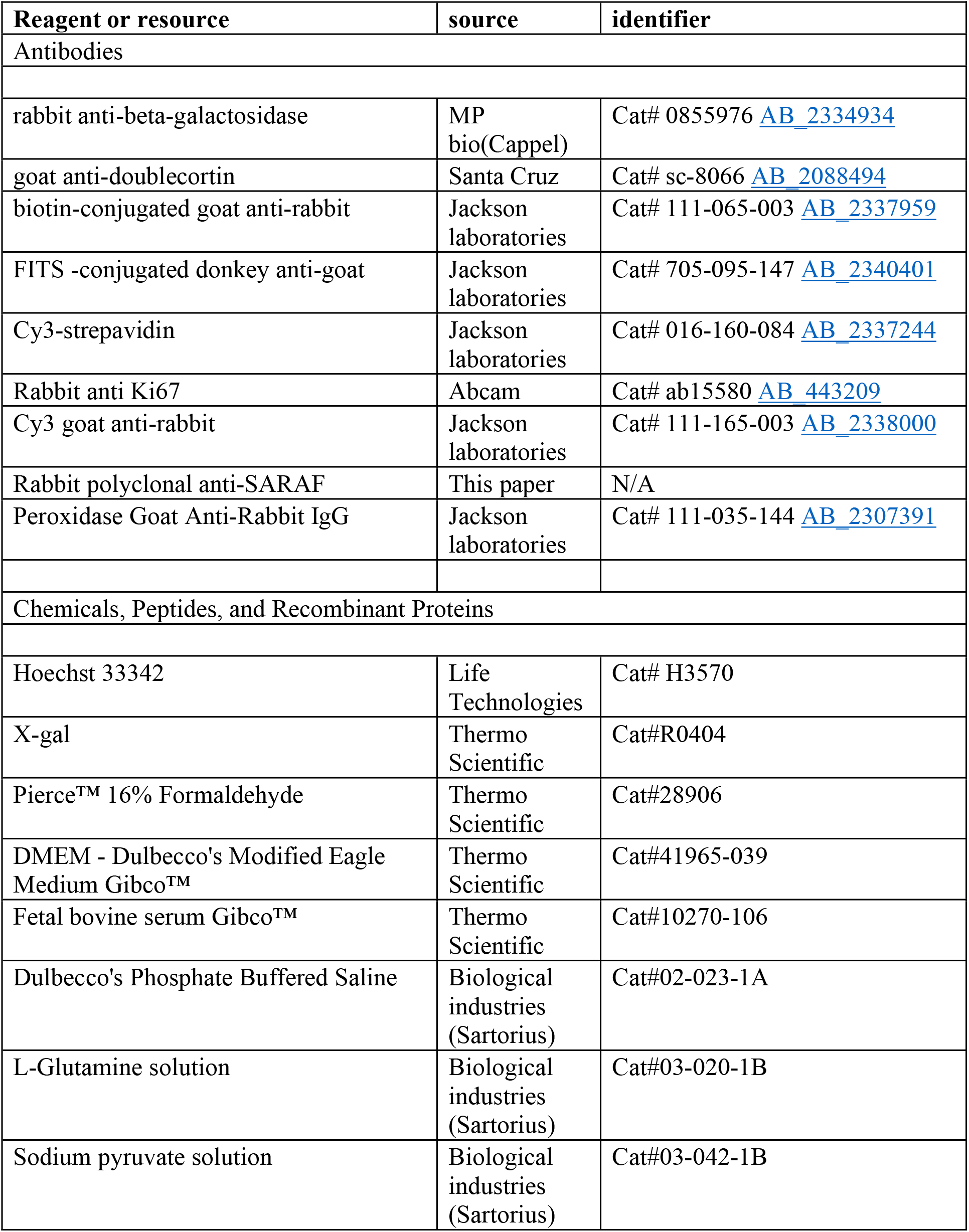
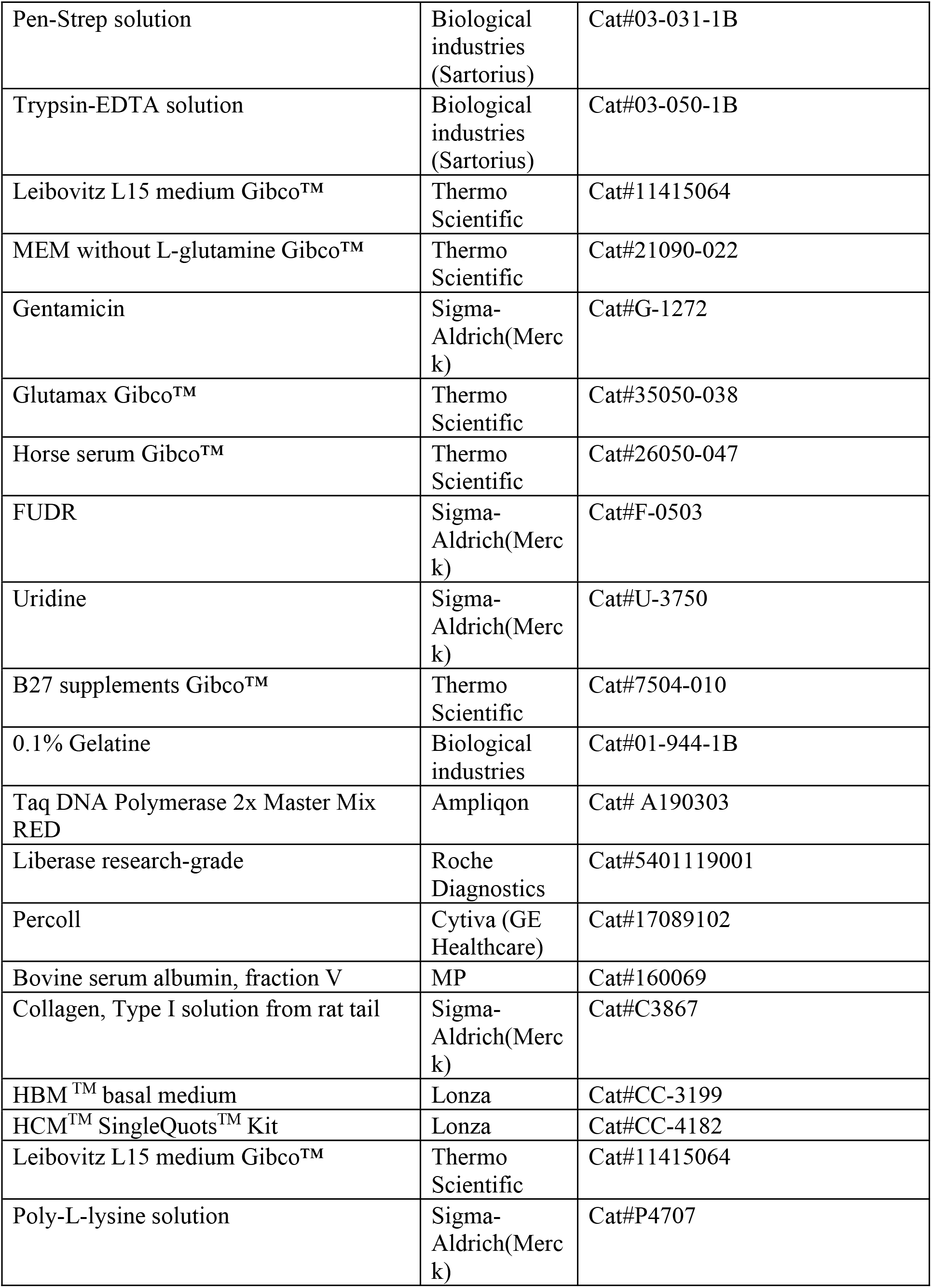
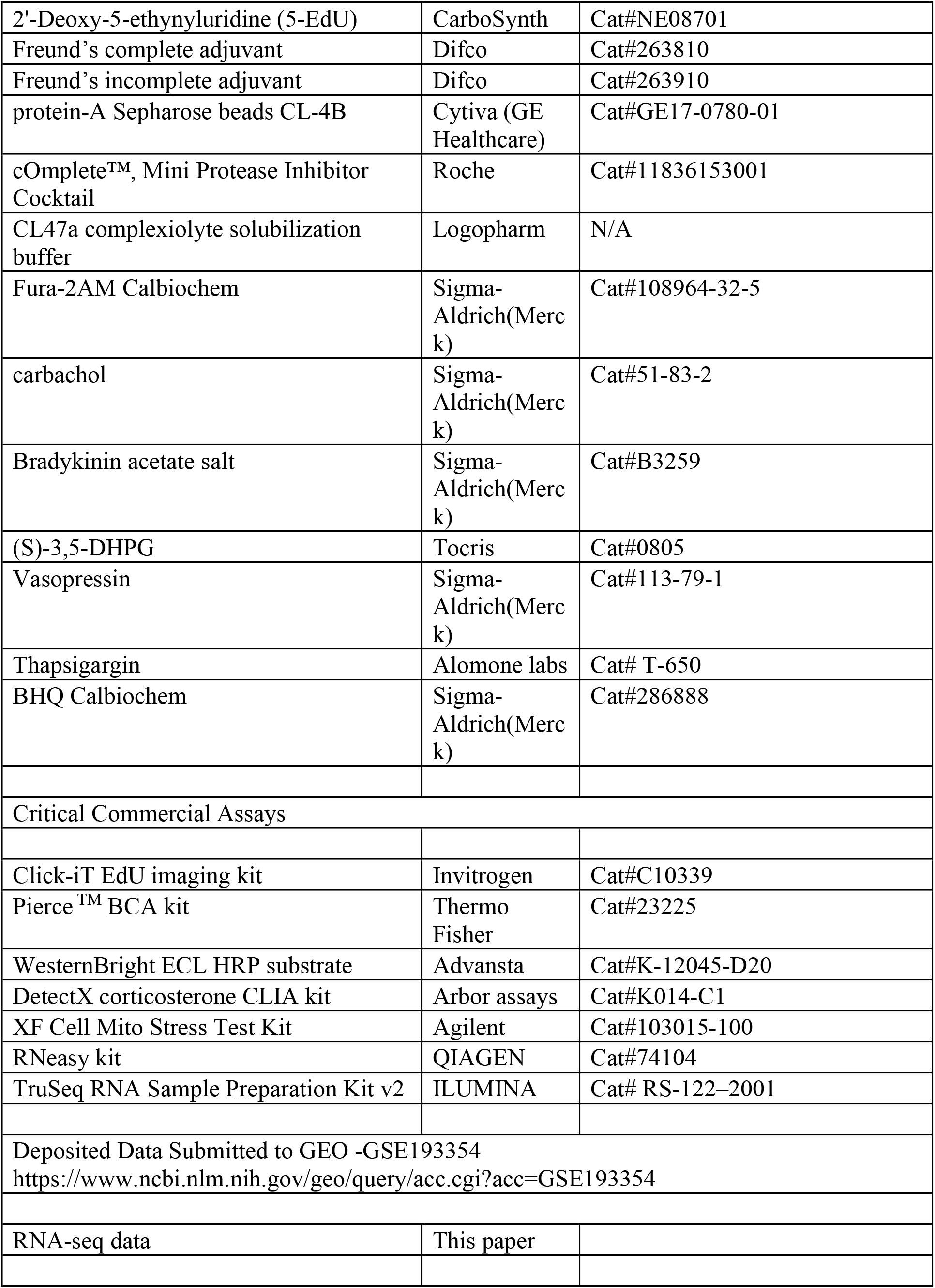
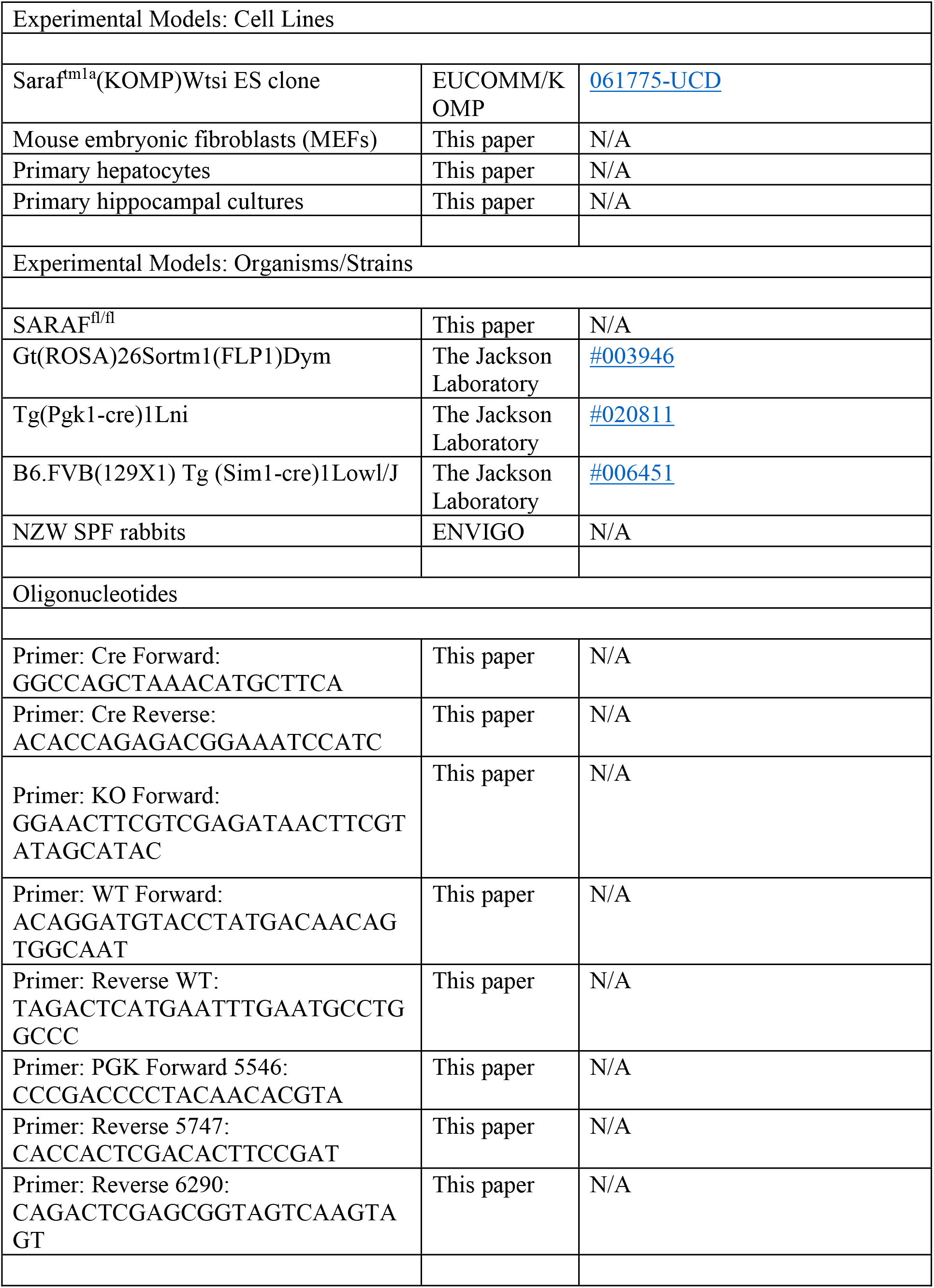
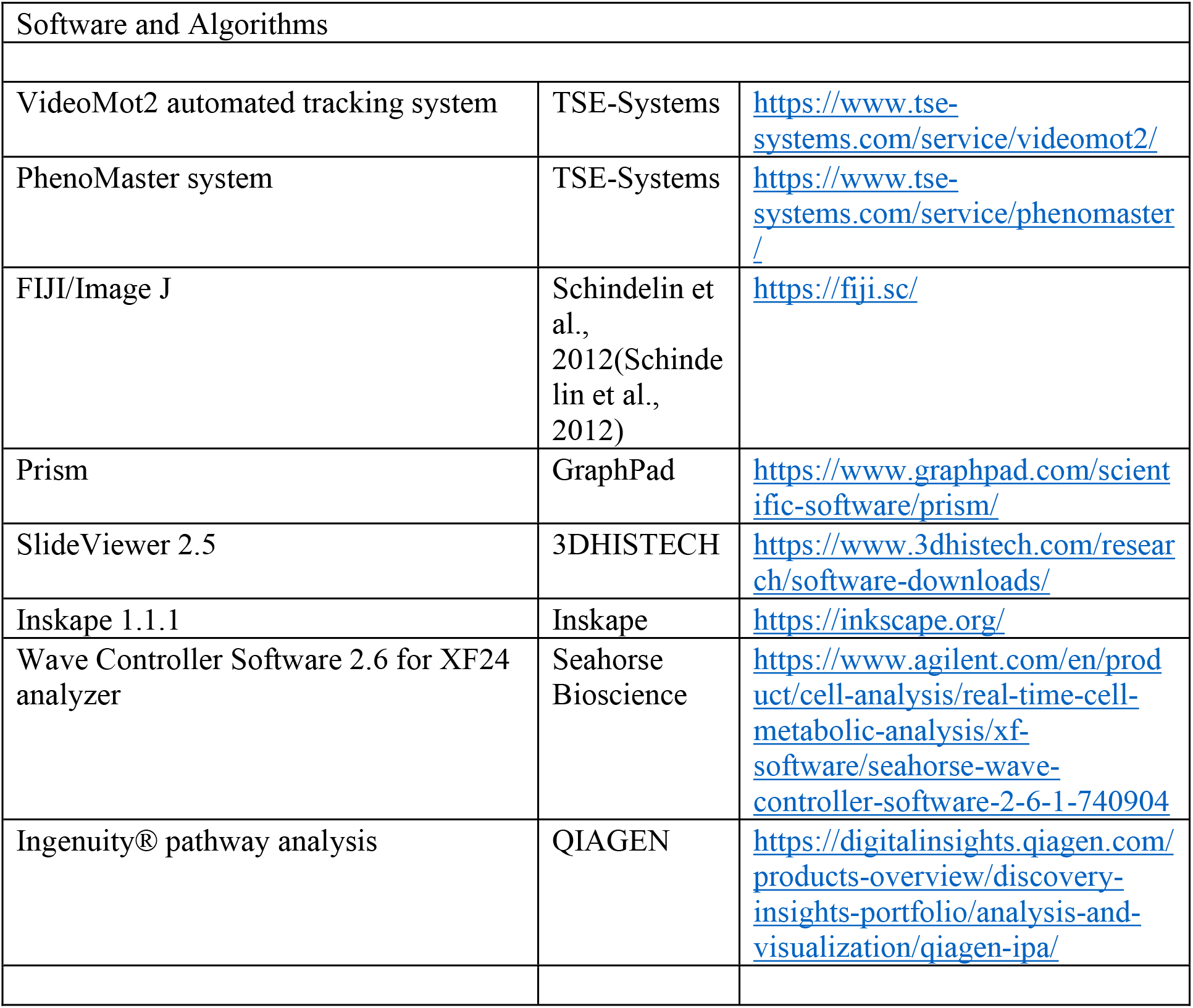

## Figure legends

**Figure 1S:**
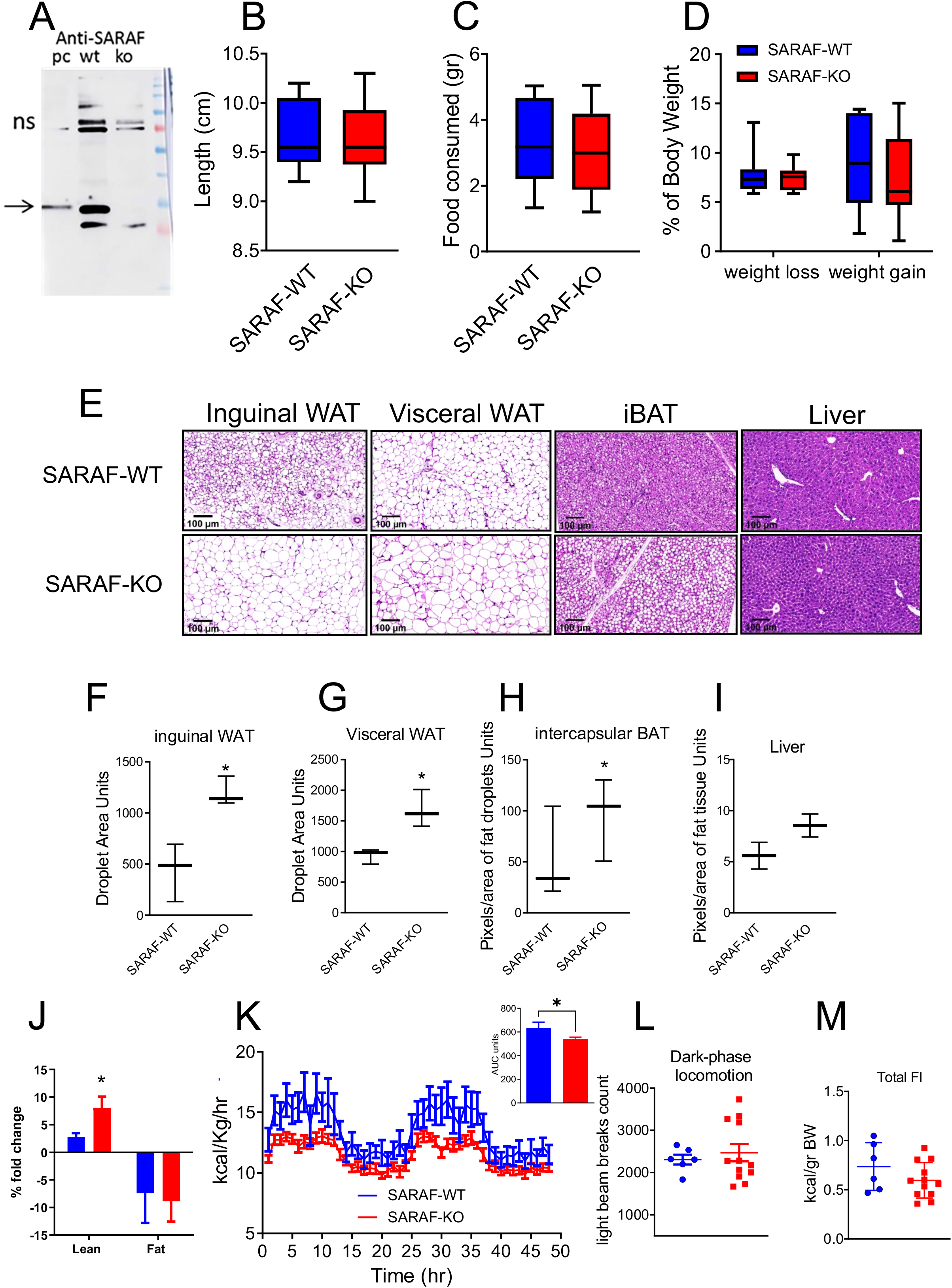
Characterization of SARAF-KO metabolic phenotype. **A**, Western blot of the whole-brain extracts from SARAF-WT, and SARAF-KO mice and from HEK293 cells (positive control, PC). **B**, Linear growth measurement of one year-old mice (SARAF-WT, n=10; SARAF-KO, n=6). **C**, food consumption 8 h refeed experiment, after 5 hours fasting during active phase of three-months-old mice (SARAF-WT, n=7; SARAF-KO, n=9). **D**, Percentage of weight loss and gain following refeed experiment of three-months old mice (SARAF-WT, n=7; SARAF-KO, n=9. **E**, histological sections of inguinal and visceral WAT, iBAT and liver from three-month-old SARAF-WT, and SARAF-KO mice. Size bar-100μm. **F-I**, quantification sections as outlined in E, Inguinal, and visceral WAT droplet size quantification and iBAT and liver pixels of fat droplets/area quantification. **J**, Four-weeks voluntary wheel training effect over fold change of lean/body weight and fat/body weight ratios. **K-M**, PhenoMaster calorimetry metabolic analysis of one year-old mice after four-week voluntary wheel training. **K**, Heat production over time (SARAF-WT, n=6; SARAF-KO, n=12). Insert AUC heat production. **L**, Dark-phase locomotion. **M**, Total food intake.

**Figure 2S:**
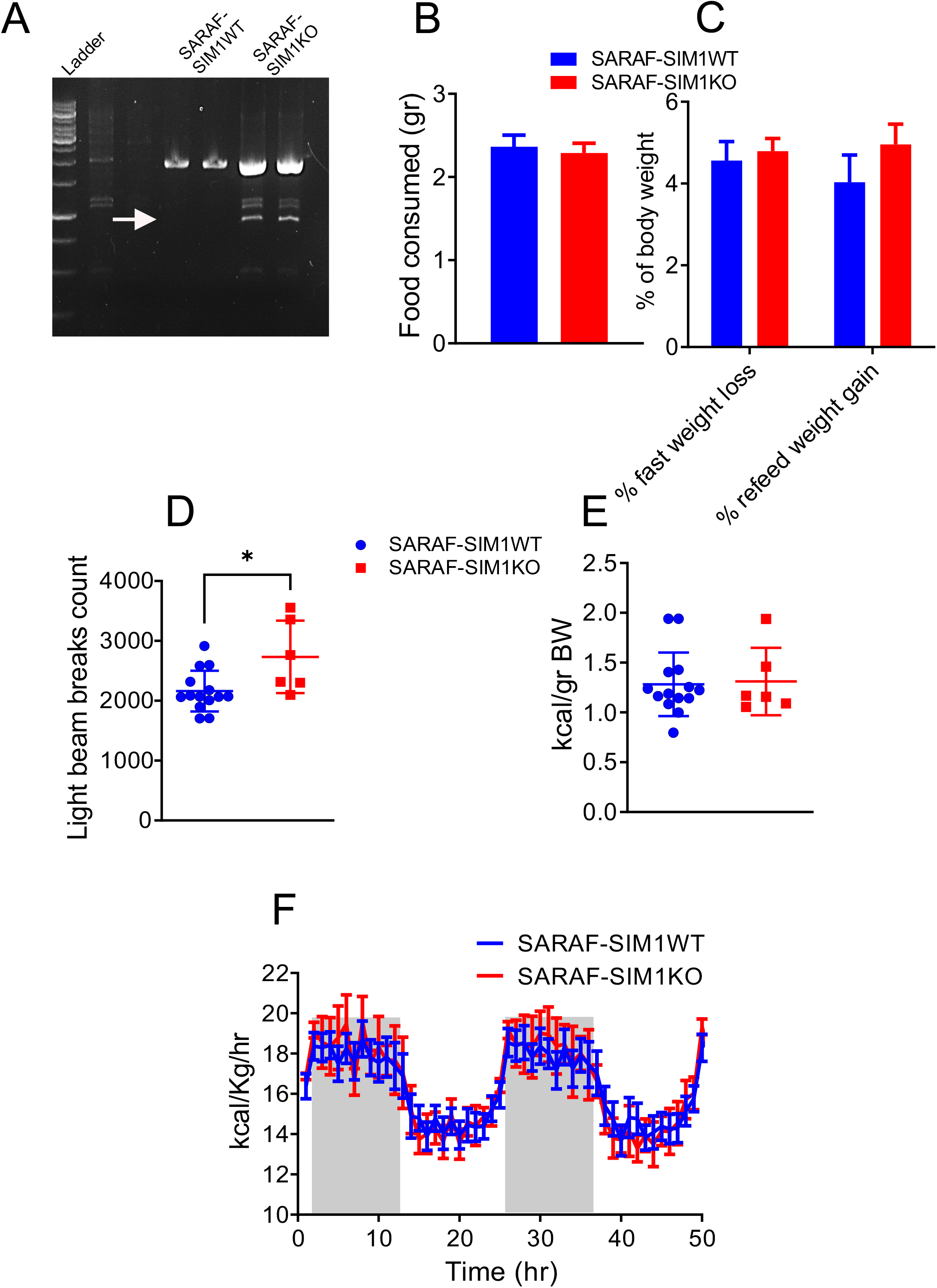
Characterization of SARAF-SIM1KO metabolic phenotype. **A**, Tissue punch PCR validation of SARAF exon excision in the hypothalamus PVN, arrow indicates the product after the removal of exon 3. **B**, Food consumed after five hours fasting at eight hours refeed experiment in 1-years-old SARAF-SIM1WT and SARAF-SIM1KO mice. **C**, percentage of weight loss after five hours fasting, and gain after eight hours refeed experiment in 1-years old SARAF-SIM1WT (n=10) and SARAF-SIM1KO (n=6) mice. **D-F**, PhenoMaster calorimetry metabolic analysis of 3-month-old SARAF-SIM1WT and SARAF-SIM1KO mice **D,** Heat production over time (SARAF-SIM1WT, n=14; SARAF-SIM1KO, n=6). **E**, Dark-phase locomotion. **F**, Total food intake.

**Figure 3S:**
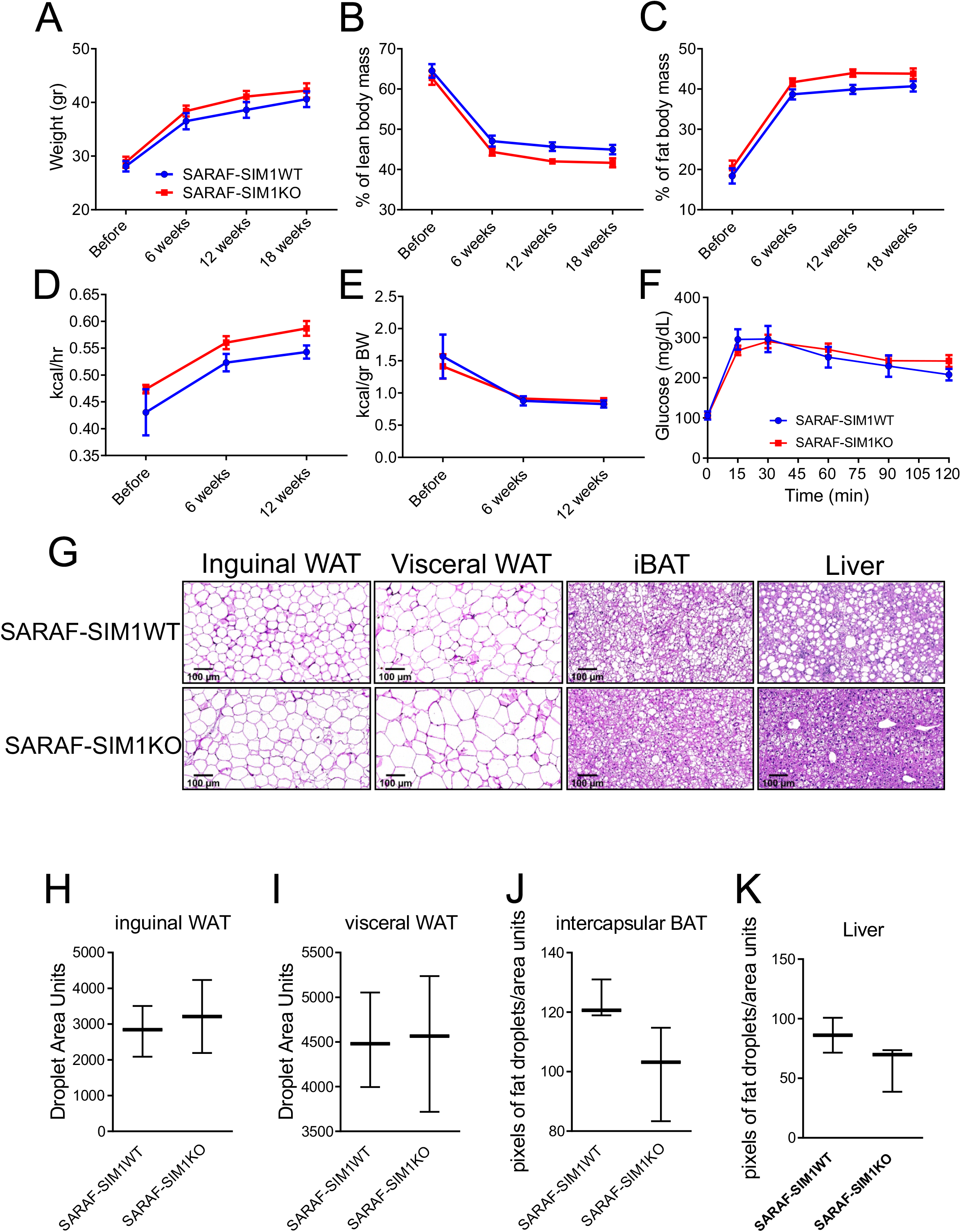
Metabolic phenotype of western diet fed SARAF-SIM1WT and SARAF-SIM1KO. **A**, mice body weight (SARAF-SIM1WT, n=5; SARAF-SIM1KO, n=11), **B**, Percent lean mass change, and **C**, Percent change in body mass over a period of eighteen-weeks on western diet. **D-E**, PhenoMaster calorimetry metabolic analysis during twelve weeks on western diet fed mice. **D**, Heat production, **E**, Total food intake. **F**, Glucose tolerance test at twelve-weeks on western diet. **G**, Inguinal and visceral WAT, BAT and liver H&E-stained tissue from mice after eighteen weeks western on fed diet. **H-K** fat droplets/area quantification of the sections as shown in G.

**Figure 4S:**
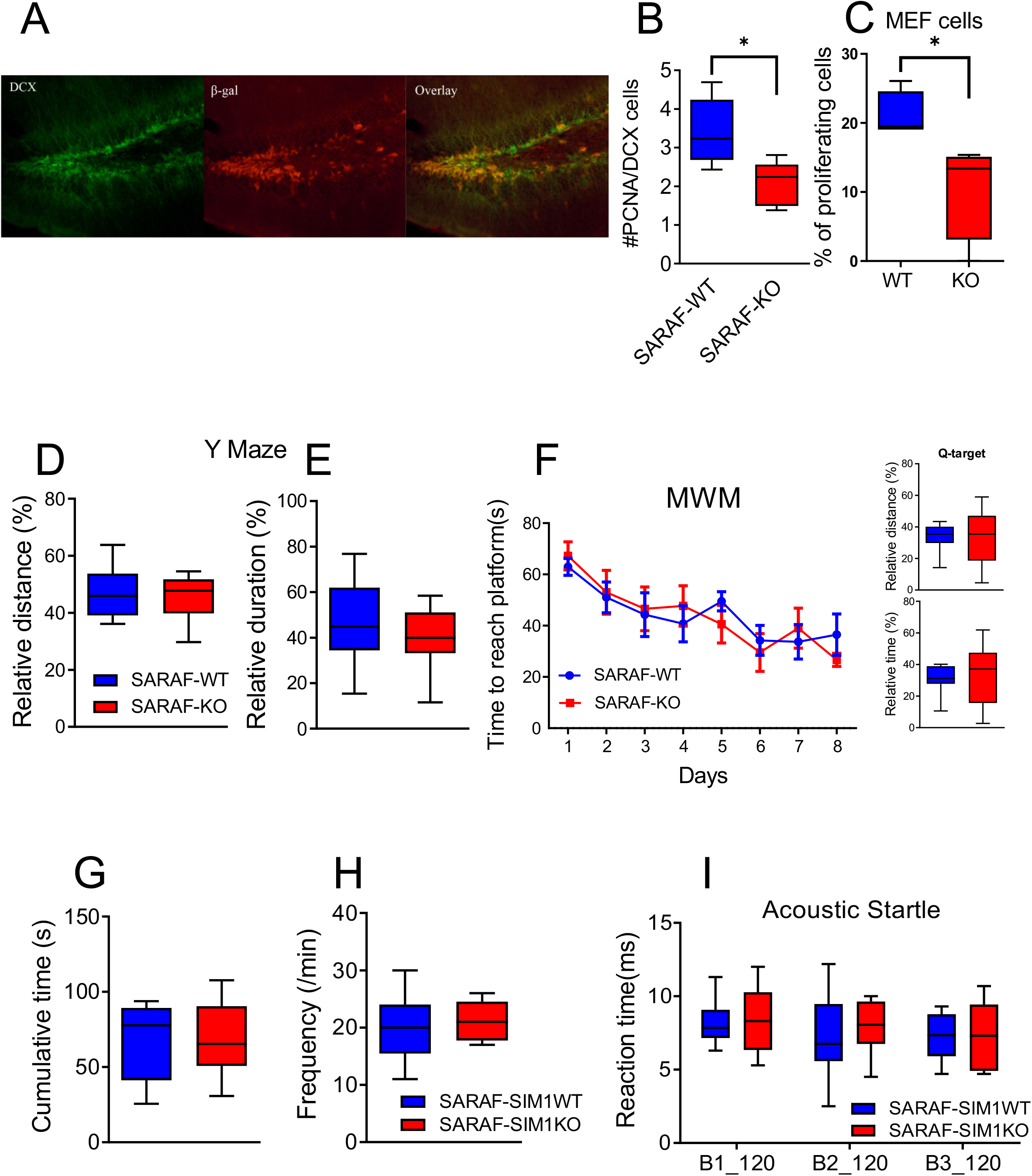
Characterization of SARAF-KO and SARAF-SIM1KO mice memory and anxiety-related phenotypes. **A**, Co-immunostaining of doublecortin (DCX) and β-gal and in hippocampal section from *saraf^m1a(KOMP)Wtsi^* line. **B**, SARAF-WT and SARAF-KO ventral hippocampal dentate gyrus proliferation quantification, number of PCNA positive cells out of DCX positive cells. **C**, Assessment of proliferation using Ki67 in MEF cells. **D-E,** Y-maze assay short memory test (SARAF-WT, n=11; SARAF-KO, n=10) and **F**, Morris Water Maze (MWM) long-term memory assay of three-months old of SARAF-WT (n=7) and SARAF-KO (n=11) mice. **G-H** DLT. G, Time in light. F. The frequency of Visits in lit section of three-months old SARAF-SIM1WT and SARAF-SIM1KO mice. **I**, Acoustic startle response tests. Reaction time in 3 blocks of 120db stimuli of three-month old SARAF-SIM1WT (n=14) and SARAF-SIM1KO (n=6) mice.

